# *Schistosoma haematobium* and *Schistosoma bovis* first generation hybrids undergo large gene expressions remodeling consistent with species compatibility

**DOI:** 10.1101/2023.05.10.540117

**Authors:** Eglantine Mathieu-Bégné, Julien Kincaid-Smith, Cristian Chaparro, Jean-François Allienne, Olivier Rey, Jérôme Boissier, Eve Toulza

**Affiliations:** Department of Environmental Sciences, Zoology, University of Basel, Vesalgasse 1, 4051, Basel, Switzerland; IHPE, Univ. Montpellier, CNRS, Ifremer, Univ. Perpignan Via Domitia, Perpignan France; CBGP, IRD, CIRAD, INRAE, Institut Agro, Univ Montpellier, Montpellier, France

**Keywords:** Hybridization, Hybrid vigor, Schistosomes, Over-dominance, Gene expression

## Abstract

When two species hybridize, the two parental genomes are brought together and some alleles might interact for the first time. To the date, the extent of the transcriptomic changes in first hybrid generations, along with their functional outcome constitute an important knowledge gap especially in non-model organisms. Here we explored the molecular and functional outcomes of hybridization in first-generation hybrid between the blood fluke parasites *S. haematobium* and *S. bovis*. To this aim we fist assembled a hybrid reference transcriptome allowing us to measure gene expression in both parental species and hybrids to further described and quantified the profiles of gene expression encountered in hybrids. We identified large-scale gene expression remodeling since up to 55% of genes were differentially expressed in hybrids compared to at least one of the parental species. We further showed that intermediate level of expression in hybrids compared to parental species was the most prevalent expression profile (38% of genes) in agreement with *S. haematobium x S. bovis* high genomic compatibility and limited divergence. Also, among the differentially expressed genes of each of the identified profiles (intermediate, under-, over-expressed, or matching the expression of one of the parental species), only a few biological processes were found enriched without patterns being consistent through crosses and sexes. Our findings suggest that hybrid heterosis could be due to change in expression of a large portion of random genes which affect various biological processes.

**Author Summary:** When two species manage to produce a viable offspring, due to different parental genetic material, new allelic interactions might arise. We do not know much about how genes are expressed in such hybrids compared to their parental species and especially for non-model species. Here we gene expressions in first generation hybrids of two blood flukes’ parasites *Schistosoma haematobium* and *S. bovis.* We quantified and categorized gene differentially expressed in first generation hybrids. We showed that more than half the genes of the hybrids were differentially expressed compared to at least one of their parental species, hence showing major gene expression remodeling occurring during hybridization. However, the most common hybrid expression pattern was intermediate level of expression compared to parental species and various but no specific biological processes were associated with hybrids differentially expressed genes. We hypothesized that because *S. haematobium* and *S. bovis* remain very compatible, when hybridizing many non-specific genes can have their expression level remodeled hence impacting the overall molecular machinery of the offspring while still sustaining highly operational hybrids.

## Introduction

Species definition has for long fueled debates among biologists due to the difficulty to draw clear boundaries between such biological entities [1]. The study of hybridization offers the possibility to shed light on the process of species formation. So far, hybridization has been mostly studied in terms of populations structure, genome structure and phenotypic outcomes [2,3]. However, the molecular mechanisms underlying the viability and performances of hybrids still constitute an important knowledge gap, especially regarding empirical evidences. To date most studies focusing on the molecular basis underlying hybrid phenotypes have been conducted either on model laboratory species or on species of economic interest [4–6]. Hence, important research avenues lay onto the gene expression, epistasis and epigenetics marks of hybrids resulting from non-model parental lines (i.e., what one could call ecologically realistic hybrids).

The study of hybridization has been mostly motivated by two extremes of hybrid phenotypes, namely hybrid incompatibilities and heterosis (or hybrid vigor, [7]). Hybrid incompatibilities describes the case when the hybrid phenotype cannot be maintained or achieved due to conflicts in the expression of each parental species genomes (i.e., *genome clash*, [7]). At the other end of the spectrum, heterosis is a phenotype characterized by higher performance of the hybrids compared to their parental lines in term of growth, size or reproductive potential [8,9]. Heterosis may provide a particular benefit in farmed species because it increases their yield, explaining why most of the literature on heterosis has focused on species of agroeconomic interest. Most first-generation hybrids are sterile or non-viable at all due to parental genome conflicts, though when viable they usually display vigor on at least some traits [7]. Interestingly, if hybrid vigor can emerge in the first generation of hybridization, this phenotype is not maintained through subsequent generations that usually show a decrease in heterozygosity (which get them closer to the parental phenotype) and may ultimately display hybrid breakdown (i.e., weaker performance than both parental strains, [10]). Hence the “hybrid phenotype” is a dynamic process throughout hybrid generations that depends on alleles that are brought together at each generations and the way they interact with each other’s [8].

Similarly, the process of hybridization does not occur equally among species since permeability toward hybridization may drastically vary between genomic backgrounds. Even after millions of years some species still display significant levels of admixture whereas other species, although close genetically, have evolved strong reproductive isolation mechanisms that prohibit hybridization either at early stages of reproduction (pre-zygotic barriers) or along the hybrid lifespan (post-zygotic barriers, for instance when hybrids die prematurely or are sterile) [11]. Once pre-zygotic barriers passed, successful hybridization then depends on parental genomes’ interaction and environmental selection. For instance, differences in the karyotype generally results in non-compatible genome interaction and non-viable offspring [12]. At the gene level, when two species hybridize, some alleles that have not been brought together might interact to result in an hybrid phenotype [13]. Depending on the distance between species and notably the difference among their transcription factors, hybrids might form and prosper whereas other genetically distant would not [13]. Some theoretical gene-level models have been developed to explain exacerbated hybrid phenotypes such as hybrid vigor (eg., dominance and over-dominance models) but so far, they remain mostly explored in species of agronomic interests and provided sometime contradictory results [13]. For species hybridizing naturally with significant introgression levels, even description of gene expression patterns remains to the date rarely documented.

Among naturally hybridizing species, a growing interest has emerged for parasites and/or their vectors since their geographic range are affected by global change (including human migration, habitat loss, global trade and climate change) hence bringing together pathogens that were previously not in contact [14]. This is particularly worrying due to the potential for exacerbated virulence in hybrid parasites [15]. Viable hybridization in parasites often results in stronger detrimental effects on their hosts, but also broader vector/host range, which carry the potential for zoonotic transmission when parasites of wild or farmed animals interact with human parasites [14]. For instance, introgression between two subspecies of *Trypanosoma brucei* have been shown to be associated with different levels of virulence [16]. Similarly, the high competence of malaria vectors, as revealed by their broad definitive host spectrum that encompassed various mammals and birds species, is thought to result from genetic admixture between *Anopheles* species which is inferred from their high levels of introgression [14,17,18]. To date most studies have focused on identifying and quantifying parasite hybrids in the wild. Although necessary to assess the risks associated with parasite hybridization, very rare studies are currently focusing on the molecular bases of the hybrid phenotypes in parasites (but see [19]).

*Schistosoma haematobium* and *S. bovis* are two blood fluke parasites for which hybrid vigor has been documented in the laboratory [20] and is suspected to have impacts on human morbidity in the field [21]. Schistosomes are the etiologic agents of Schistosomiasis (or Bilharziasis) a Neglected Tropical Disease affecting over 240 million people in the world [22]. These gonochoric parasites have a two-host life cycle including a mollusc intermediate host in which asexual multiplication occurs, and a mammalian definitive host in which sexual reproduction (separate sexes) takes place. In the latter host, parasite species co-infection can lead to interbreeding and the production of viable hybrid progeny [23,24]. Several field or experimental viable interspecific crosses have been evidenced among the *Schistosoma* genus [25]. Depending on the phylogenetic distance between the interacting species, the genetic background of the resulting progeny varies between parthenogenic (hence non-viable introgressed individuals) to fully viable introgressed individuals [26]. *Schistosoma haematobium* and *S. bovis* are sister species amongst the *haematobium* clade [27]. *Schistosoma haematobium* is specialized toward humans as definitive hosts, while *S. bovis* is a livestock and rodent parasite. However, despite apparent host-specific barriers, hybrids between *S. haematobium* and *S. bovis* are observed in the field with common occurrences in several African countries including Senegal, Benin, Mali, Niger, Cameroon, Ivory Coast or Nigeria [28–35]. *Schistosoma haematobium x S. bovis* hybrids were also involved in the European outbreak that occurred in Corsica in 2013 and where infections are still persisting and expanding to nearby rivers [36,37]. Both for human health impacts and epidemiological aspects, schistosome hybrids present worrying issues. Since schistosome first-generations hybrids tend to produce more eggs with larger size than parental forms [25,38–40] and because the presence of eggs trapped in human tissues are responsible for the physiopathology of the disease, hybridization among schistosomes is expected to affect the parasites induced pathology and transmission with important consequences for public health and disease control.

In this study we aimed at characterizing gene expression levels and molecular pathways observed in *S. haematobium* and *S. bovis* first generation hybrids. Relying on a genome-guided transcriptomic approach we first assembled a common reference transcriptome for *S. haematobium, S. bovis* and their hybrids to compare their gene expression levels. We categorized hybrid expression profiles by comparing them to both parental lines. Hence, we distinguished five expression profiles of the hybrids namely: over-expressed, under-expressed, intermediate (also referred as additive expression, [41,42], *S. haematobium-*like and *S. bovis-* like expression profiles. Finally, among genes belonging to each category of expression profiles we tested whether some biological processes were significantly enriched (i.e. over-represented). Based on current pieces of evidences on heterosis built upon cultivated species and model organisms, we could expect to observe most genes having a non-intermediate profile (over-, under-expressed and parent-like) that may sustain an increased expression of hybrid vigour traits (such as reproduction or growth) when comparing the hybrids to at least one of the parental species. However, expression profiles may depend on the overall parental genome’s compatibility: the more two genomes are compatible, the less conflict in expression could be expected and the more likely genes would have an intermediate expression profile.

## Material and Methods

### Ethics approval

Housing, feeding and animal care, including experiments on animals were carried out according to the national ethical standards established in the writ of 1 February 2013 (NOR: AGRG1238753A) setting the conditions for approval, planning and operation of establishments, breeders and suppliers of animals used for scientific purposes and controls. The experiments carried out for this study were approved and provided a permit A66040 for animal experimentation by the French Ministry of Agriculture and Fishery (Ministere de l’Agriculture et de la Peche), and the French Ministry for Higher Education, Research and Technology (Ministere de l’Education Nationale de la Recherché et de la Technologie). The investigator has the official certificate for animal experimentation, obtained from both ministries (Decret n^◦^ 87/848 du 19 octobre 1987; authorization number 007083).

### Study model

#### Parasite parental species and reciprocal experimental crosses

*Schistosoma haematobium* was initially recovered from infected patients in the Southeast part of Cameroon (Barombi Kotto lake, 4°28’04”N. 9°15’02”W) in 2015. Specifically, eggs from positive urine samples were collected and we exposed local intermediate hosts (*Bulinus truncatus*) to five miracidia before they were transferred to the laboratory. Parasites were further maintained through completion of their cycle using Golden hamsters (*Mesocricetus auratus*) as definitive hosts and *B. truncatus* as intermediate host [24]. *Schistosoma bovis* was recovered from the Spanish laboratory of parasitology of the Institute of Natural Resources and Agrobiology in Salamanca, and maintained in the laboratory with both *Bulinus truncatus* and *Planorbarius metidjensis* sympatric intermediate hosts and on *M. auratus* definitive hosts [43]. Starting with experimental crosses, we first created a full-sibling consanguine line of *S. haematobium* because of its potential high genetic diversity compared to the *S. bovis* isolate maintained in laboratory for several decades. To do so we crossed a single male with a single female after molecular sexing of the parasite larvae [44]. Briefly, batches of *B. truncatus* were individually infected with a single miracidium of *S. haematobium*, allowing the development of clonal male or female genotype. At patency (i.e., 55 days after exposure) *cercariae* genotypes emitted from individual molluscs were sexed and parasites from two molluscs (one infected with a male genotype and one with a female genotype) were used to infect hamsters (300 cercariae of each sex) by surface application method for one hour. The resulting offspring constituted our full-sibling consanguine line used for subsequent experiments. Methods employed for mollusc and rodent infections and parasite collection were described previously [45–47].

To recover parental species adult worms and assess their gene expression profiles, we infected six hamsters with homospecific combinations of 600 cercariae (300 cercariae of each sex) of *S. haematobium*, or *S. bovis*. Briefly, for both parental species, a pool of *miracidia* obtained from homospecific crosses was used to infect molluscs (5 *miracidia* per mollusc). Fifty-five days after exposing the molluscs to the parasites, pools of 600 *cercariae* from infected molluscs were used to infect hamsters. Three months after infection hamsters were euthanatized and adult worms were recovered by hepatic perfusion.

Laboratory *S. haematobium x S. bovis* hybrids were produced by exposing molluscs (*B. truncatus* or *P. metidjensis*, respectively) to a single miracidium of either parasite species and using the resulting single sex clonal pools of *cercariae* of each species to infect hamsters with mixed combinations of parasites in equal proportions (Fig 1). *Schistosoma haematobium* and *S. bovis* were reciprocally crossed to produce F1 (female *S. haematobium* x male *S. bovis*) and F1’ (female *S. bovis* x male *S. haematobium*) hybrids. Batches of six hamsters per combinations were infected using 300 female *S. haematobium* and 300 male *S. bovis cercariae* (F1 cross; n=6) or 300 male *S. haematobium* and 300 female *S. bovis cercariae* (F1’ cross; n=6). Three months and twenty days after infection, hamsters containing the adult parents (heterospecific couples) and hybrid egg progenies were euthanized.

**Fig 1:**
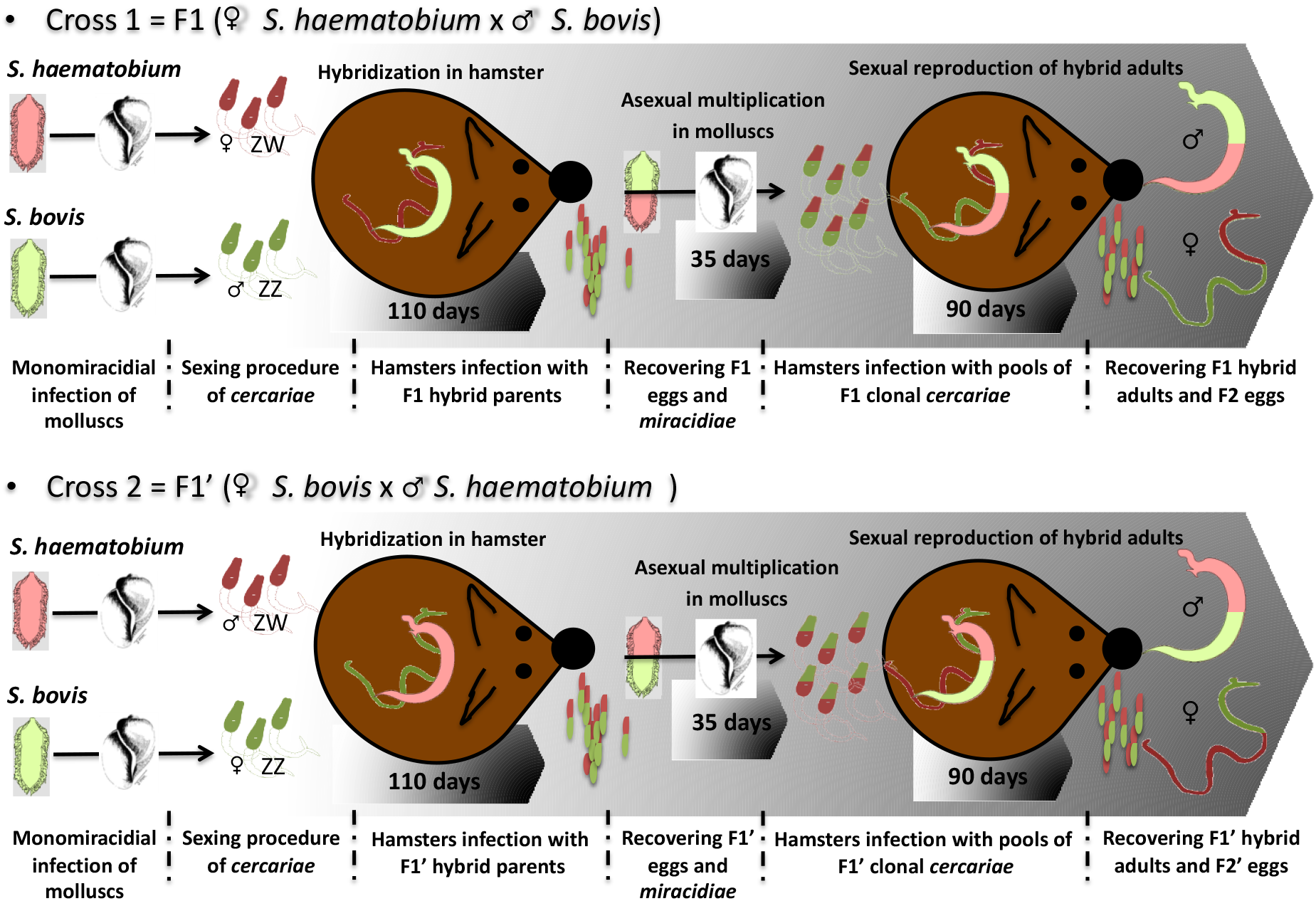
Experimental infections to produce *S. haematobium x S. bovis* reciprocal first-generation hybrids.

In order to recover first generation reciprocal F1 and F1’ hybrid adult worms and evaluate their expression profiles, hybrid eggs were hatched and hybrid *miracidia* were used to infect *B. truncatus* snails (n=5 *miracidia* per mollusc). After the prepatent phase (∼35 days) the F1 and F1’ *cercariae* were used to infect hamsters (n = 7 hamsters for each cross) with pools of 600 cercariae of unknown sex, and F1 and F1’ adult worms were collected from portal perfusion at 90 days after infection. A graphical presentation of the experimental cross design is presented in Fig. 1

#### RNA extraction and transcriptome sequencing procedure

Using a binocular microscope and a small paintbrush we separated the paired adult worms according to their sex to avoid any contamination from the other sex. We constituted pools of 10-12 individuals per condition (*S. haematobium, S. bovis,* F1 and F1’) and sex in three biological replicates representing a total of 24 samples (4 conditions x 2 sexes x 3 replicates). These samples were directly flash frozen in liquid nitrogen, and stored at -80°C. Total RNA extractions were performed using the TRIzol Thermo Fisher Scientific protocol (ref: 15596018) slightly modified as each reagents volume was halved. Briefly, pools of frozen adult worms in 2ml microtubes were grounded with two steel balls using Retsch MM400 cryobrush (2 pulses at 300Hz for 15s). After extraction following manufacturer’s protocol, total RNA was eluted in 44 μl of ultrapure water. DNase treatment was then performed using Thermofisher Scientific Turbo DNA-free kit TM (ref: AM1907). RNA was then purified using the Qiagen RNeasy mini kit (ref: 74104) using manufacturer’s protocol and eluted in 42 μl of ultrapure water. Quality and concentration were assessed by spectrophotometry with the Agilent 2100 Bioanalyzer system and using the Agilent RNA 6000 nano kit (ref 5067-1511). Library construction and sequencing were performed at the Génome Quebec platform. The TruSeq stranded mRNA library construction kit (Illumina Inc. USA) was used following the manufacturer’s protocol with 300 ng of total RNA per sample. Sequencing of the 24 samples was performed in 2×100 bp paired-end on a Illumina HiSeq 4000. Sequencing data are available at the NCBI-SRA under the BioProject accession number PRJNA491632.

#### Transcriptome assembly and generation of a common reference between S. haematobium and S. bovis

Because parental species may share common genes, and because the hybrids are composed of alleles from both species, a single reference was needed to compare gene expression levels between parents and hybrids. We produced a non-redundant transcriptome of *S. haematobium* and *S. bovis* genes as common reference. Precisely, we aimed to distinguish between, *S. haematobium* specific genes, *S. bovis* specific genes and orthologous genes between the two species. We relied on the two most up to date genome assemblies of *S. haematobium* and *S. bovis* [40,48], and RNA-sequencing data of the two parasites species.

First, we conducted reference-based transcriptome assemblies for *S. haematobium* and *S. bovis* respectively. This transcriptome assembly was conducted using the Galaxy instance of the IHPE laboratory and of the Ifremer [49,50]. After ensuring read quality using FastQC (https://www.bioinformatics.babraham.ac.uk/projects/fastqc), we mapped *S. haematobium* and *S. bovis* paired end reads respectively on their reference genomes using the fast and sensitive mapper HISAT2 (Galaxy version 2.1.0, [51]) with default parameters. Exon/intron structure was then recovered using StringTie (Galaxy version 2.1.4, [52]) and with genomes’ annotations as extra input. The results from each sample were then merged in one single file per species (one for *S. haematobium* and one for *S. bovis*) using the tool StringTieMerge. From the resulting gtf files, spliced exons for each transcript were extracted from the two reference genomes using *gffreads* tool (Galaxy version 2.2.1.1 [53]). This step resulted in two species-specific transcriptome on which we further selected transcripts containing an open reading frame (hereafter referred as genes) with TransDeCoder (Galaxy Version 3.0.1 [54]). Then, for each transcriptome, genes were clustered based on similarity (threshold put at 96% for *S. haematobium* and *S. bovis, i.e.,* which allowed to keep redundancy percentage measured on Busco (Galaxy Version 4.0.4 [55]) under 10%) to retain only non-redundant protein coding genes. This resulted in a final transcriptome composed of respectively 11 161 genes and 10 768 genes for *S. haematobium* and *S. bovis*

From these two references we built one single reference, where *S. haematobium* genes, *S. bovis* genes and orthologous genes were indicated. To do so, we identified strict orthologous genes using a reciprocal blast hit approach. We ran a reciprocal tblastx (version 2.6.0, [56]) between *S. haematobium* and *S. bovis* transcriptomes. We set the maximum sequence target to one for each match. From blast results, we further computed coverage and global percentage of similarity over the sequence (based on the percentage of identity, query start and end, and sequence length, see online resources for the script). We finally selected best reciprocal matches based on the match having the highest global similarity and query coverage.

Finally we assessed the overall quality of our reference transcriptome in terms of representativeness and completeness [57]. Representativeness was measured as the percentage of raw reads of each species and hybrid crosses mapping back on the transcriptome and was assessed with the aligner bowtie2 in *very-fast* mode [58]. Assembly completeness was measured with Busco version 4.1.4 as the number of metazoan orthologous genes (metazoa_odb9 dataset) retrieved in the transcriptome.

#### Quantification of gene expression and differential expression analysis

The reference transcriptome we assembled was composed of both *S. haematobium* and *S. bovis* transcripts. For this reason, we used *de novo* transcriptome assembly tools to estimate gene expression levels of both parental lines and hybrids. We relied on the two Trinity tools, namely “*Align reads and estimate abundance”* and “*Build expression matrix”* to estimate gene raw counts [54]. The raw counts matrix resulted in a set of 17,191 quantified transcripts and was processed in R (version 4.0.0 [59]) using the package DESeq2 (version 1.28.1 [60]). Transcripts with less than 10 counts across all samples were excluded from further analysis hence resulting in a set of 16,613 analyzed transcripts.

We relied on the R package DESeq2, that internally corrects for library size, to calculate differential gene expression and hence to determine gene expression profiles. Based on expression profiles we defined five categories of hybrid genes that were captured setting proper contrasts in DESeq2. The *under-expressed profile* refers to genes that were under-expressed in hybrids compared to both *S. haematobium* and *S. bovis* with a significance level defined at a False Discovery Rate of 5% (FDR<0.05). The *over-expressed profile* describes genes that are significantly over-expressed in hybrids compared to both *S. haematobium* and *S. bovis* (FDR<0.05). The *intermediate profile* encompasses genes that have an expression level between those of the parents, i.e. genes that are significantly over-expressed in hybrids compared to one of the parental species while being significantly under-expressed compared to the other (FDR<0.05). The *haematobium-like profile* describes genes that are significantly differentially expressed between *S. haematobium and S. bovis* (FDR<0.05 and Log2Fold Change (LFC)>1) but that are significantly similarly expressed between hybrids and *S. haematobium* (FDR>80% and Log Fold Change (LFC)<0.05). Finally, the *bovis-like profile* refers to genes that are significantly differentially expressed between *S. haematobium and S. bovis* but that are similarly expressed between hybrids and *S. bovis.* Genes that fall into none of those gene expression profiles are genes that are not differentially expressed in hybrids and both parental lines. To capture cross-specific and sex-specific gene expression patterns, hybrids expression profiles were quantified within each sex and each cross (F1 or F1’).

#### Transcriptome annotation and functional analysis

Annotation was conducted with PLAST (version 2.3.3 [61]) on the non-redundant protein database of NCBI (released of the 2022-01-07 [62]). The transcriptome was cut in 35 chunks of 500 sequence length to optimize computing time. A maximum of twenty hits were retained per query with one single best High-scoring Segment Pair (HSP) (i.e., local alignment with no gaps that achieves one of the highest alignment scores in a given search, (Fassler and Cooper 2008.) and an e-value threshold set to 1^e-3^. Results were then loaded in Blast2Go (Omicsbox version 1.4, (Conesa et al. 2005)) to run Interproscan and pursue mapping and annotation. To test whether some Gene Ontology (GO) terms (biological processes, molecular functions or cellular component) were significantly enriched in DEG, we conducted Fisher exact test in Blast2GO (which compare within each GO the number of gene from a define gene set and to the number of genes from the reference to test whether this GO contains significantly more genes from the gene set compare to genes from the reference) followed by a False Discovery Rate (FDR) analysis to correct for multiple comparisons. To ease result interpretation, we further selected non-redundant GO terms falling into biological processes and displayed Tree Map graphical representations using Revigo with list size set to small [65]. Analyses were conducted on gene sets of each hybrid expression profile categories (under- and over-expressed, intermediate, *S. haematobium* like and *S. bovis* like) independently for each hybrid cross (F1 and F1’) and each sex (female and male).

## Results

### Reference transcriptome assembly

We were able to assemble a common reference transcriptome for *S. haematobium* and *S. bovis* that was made up of 17,191 non-redundant transcripts. In this transcriptome, 4,738 genes were identified as strict orthologous genes, 6,423 as *S. haematobium* genes, and 6,030 as *S. bovis* genes. When mapping back read sets of each species, we obtained a mapping rate of 60.35% (± 3.28) for *S. bovis* reads, of 63.42% (± 0.86) for *S. haematobium*, 64.33% (± 1.82) for F1 first generation hybrids and 64.86 (±2.23) for F1’ first generation hybrids (see Supplementary file S1: Table S1). Also, 71.5% of 954 metazoan orthologous genes were retrieved including 50.6% of those genes found in one single occurrence, and 20.9% found at least twice.

### Gene expression profiles

A Principal Component Analysis conducted on parent and hybrid samples revealed that one F1 replicate suspiciously clustered apart from others and was thus excluded from further analysis (Supplementary file S1: Fig. S1). Otherwise, as expected, F1 samples were clustering between *S. haematobium* and *S. bovis* samples (Fig S1). We categorized five typical gene expression profiles being, *over-expressed*, *under-expressed, intermediate, S. haematobium-like* or *S. bovis-like* (see Supplementary file S1: Table S1, Fig. 2, Supplementary file S1: Fig. S2-S4). After removing 578 lowly expressed genes (i.e., transcripts with less than 10 counts across all samples) from the initial 17,191 genes of the transcriptome, 16,613 genes were considered in the gene expression analysis. 9 161 (55%) were assigned to a hybrid expression profile (i.e., under-, over-expressed, intermediate or like one or the other parental species) and 7 453 (45%) genes were not assigned to a particular hybrid expression profile (those genes were similarly expressed in hybrids and both parental lines, see Supplementary file S2). The most abundant expression profile was the intermediate profile (i.e., 6 308 genes, 38%) whereas the less abundant profiles were the *S. haematobium-*like profile, the *S. bovis-*like profile and the over-expressed profiles (439, 443 and 502 genes respectively, 3% each, Fig. 2, Supplementary file S2). Also, very few genes falling into each expression profile were common between crosses (i.e., between F1 and F1’) and between sexes (i.e., between males and females, see Supplementary file S1: Fig. S5). We found common genes between crosses and sexes only for intermediate profiles (22.1% of the genes having an intermediate profile, see Supplementary file S1: Fig. S5) and under-expressed profile with 0.6% of shared genes for this profile (see Supplementary file S1: Fig. S5).

**Fig 2:**
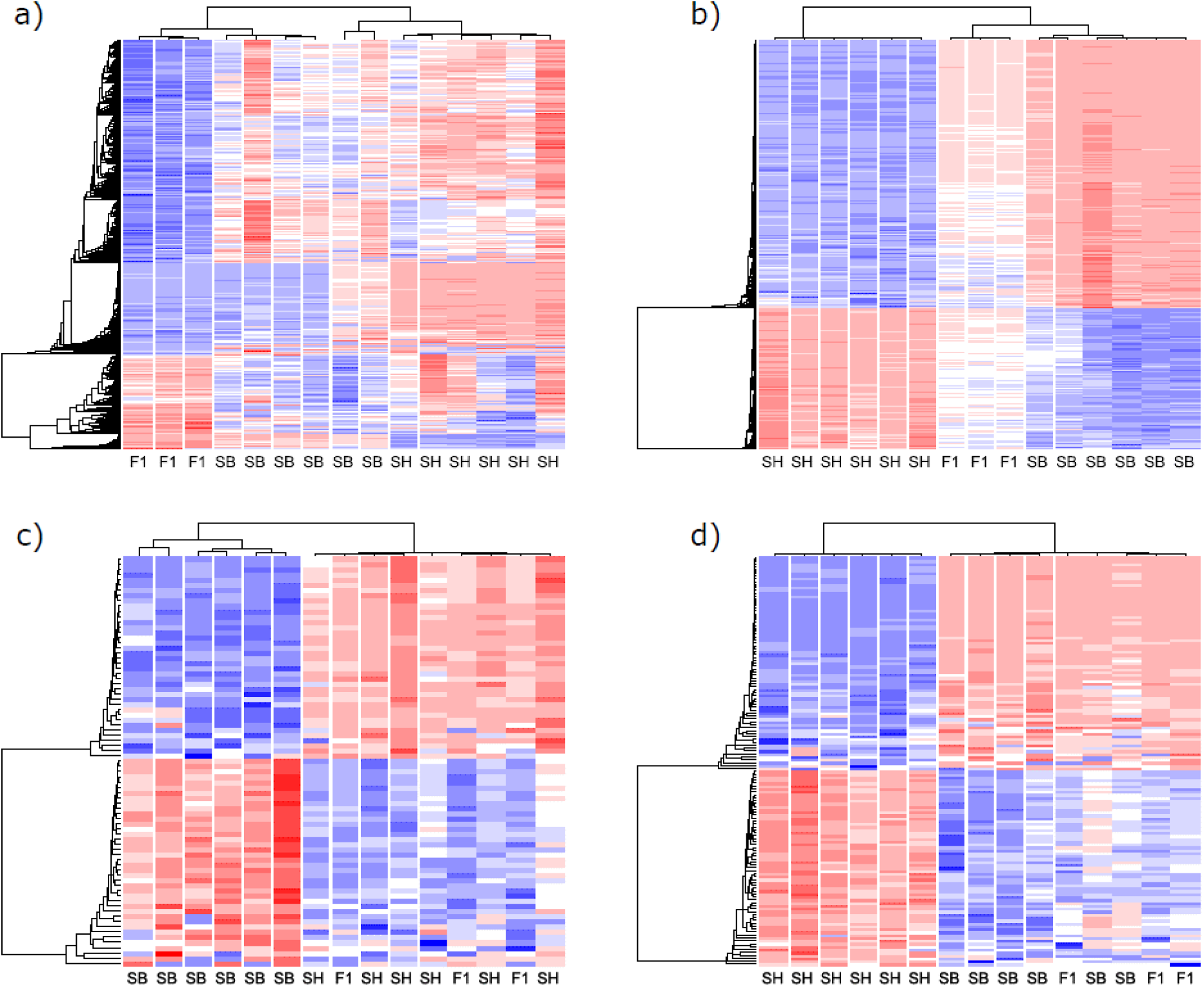
Heatmaps illustrating the gene clustering within each hybrid expression profiles in F1 females (F1) compared to *S. haematobium* (SH) and *S. bovis* (SB) females: under and over-expressed profiles (a), intermediate profiles (b), *S. haematobium* like profile (c) and *S. bovis* like profile (d).

**Fig 2:**
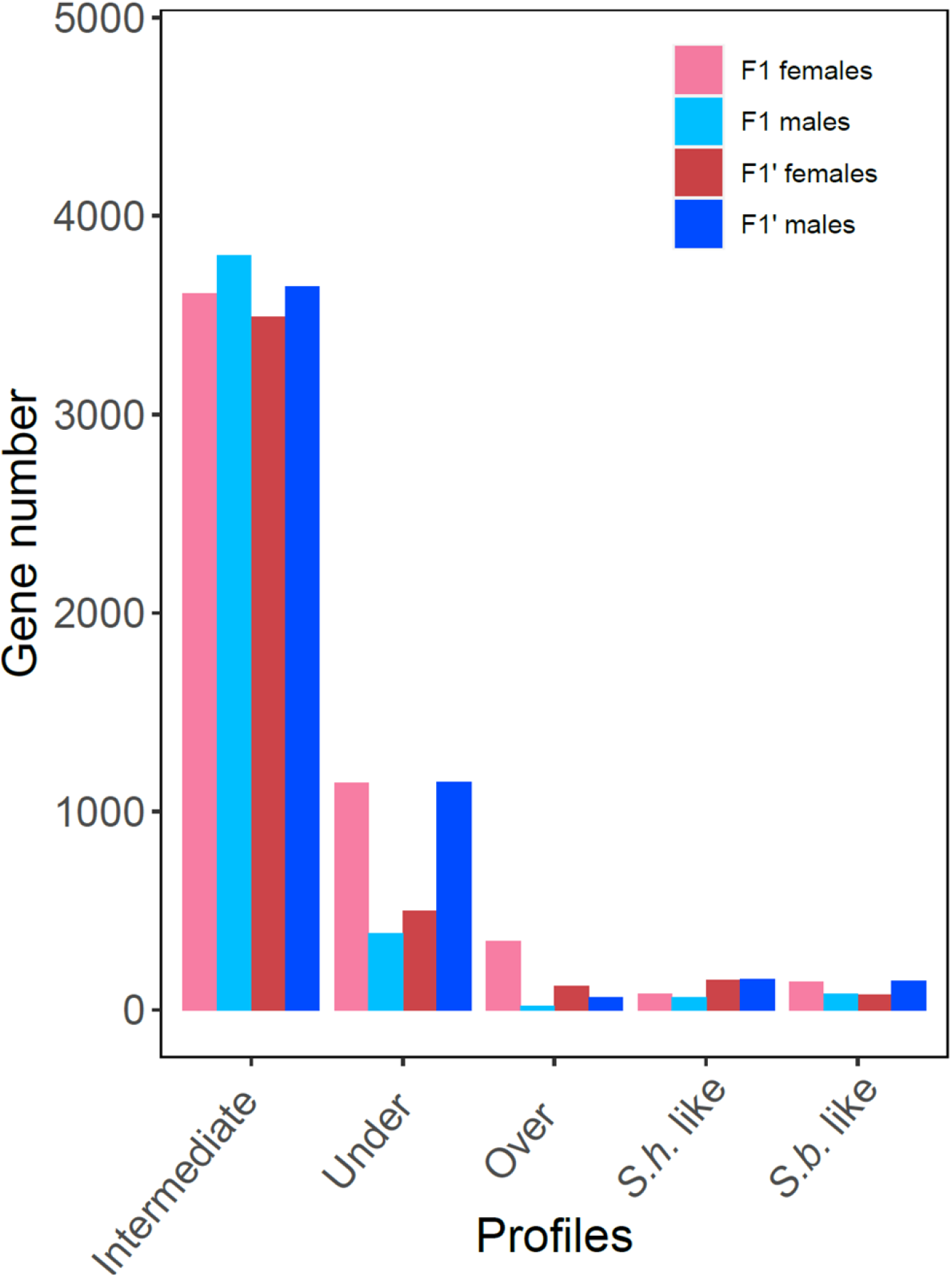
Barplot showing for males and females of each hybrid cross, the number of genes falling into the over-expressed, under-expressed, intermediate, *S. haematobium* (*S.h.*) and *S. bovis* (*S.b.*) like expression profiles.

### Functional analysis

Gene Ontology terms were found enriched in genes belonging to the five sets of gene expression profiles, namely, intermediate expression profile in F1’ males, over-expressed profile in F1 as well as F1’ females, under-expressed profile in F1’ males as well as F1 females (see Supplementary file S3, Fig. 4). No GO were found enriched with gene sets corresponding to other expressions profiles (*S. haematobium* like, *S. bovis* like, intermediate expression profiles for F1’ females, F1 and F1’males, over-expressed profiles in males and under-expressed profiles in F1’ females and in F1 males).

**Fig 4:**
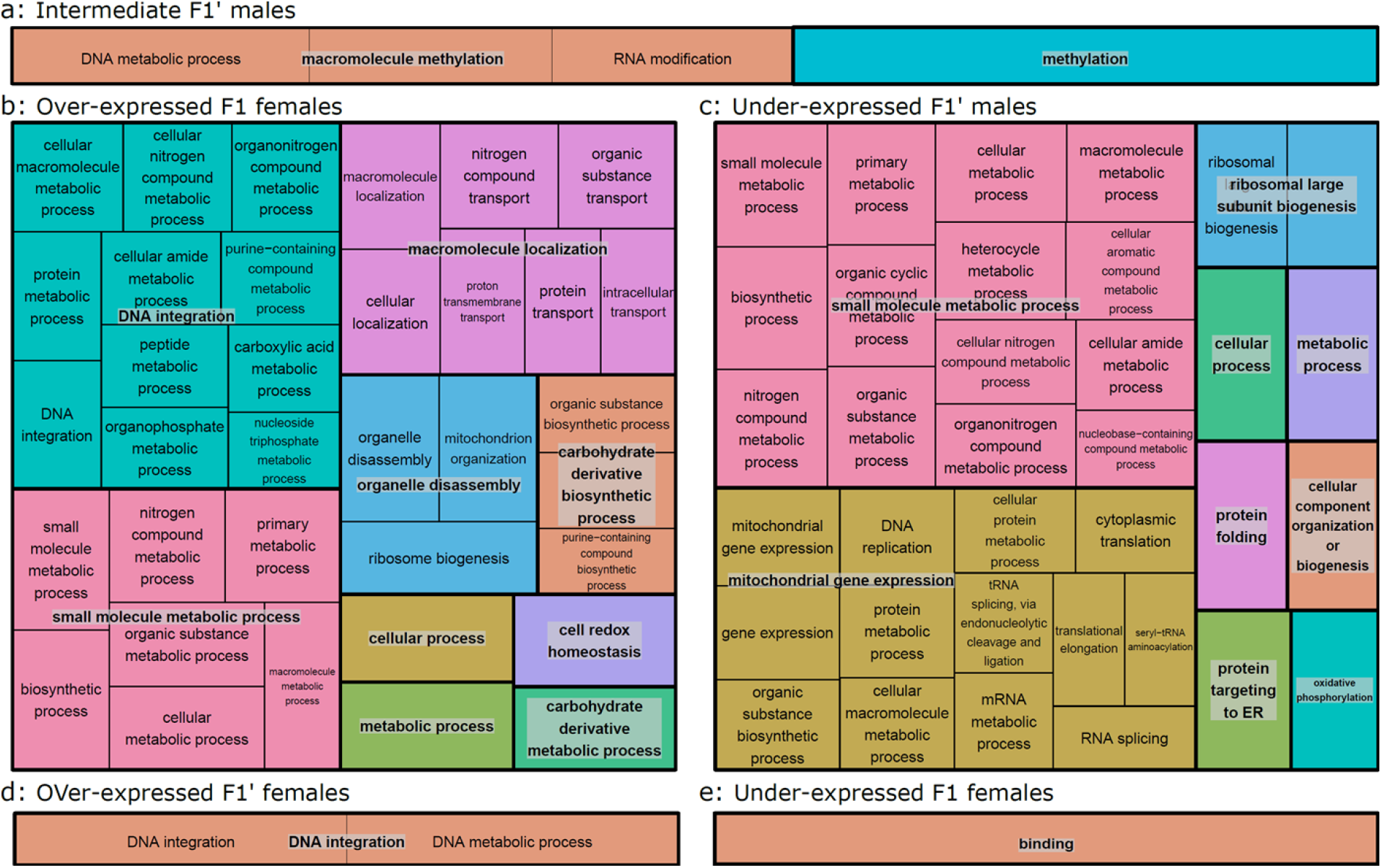
Tree maps of non-redundant biological processes enriched in genes having an intermediate profile of expression in F1 males (a), in genes having an over-expressed expression profile in F1 females (b), in genes having an under-expressed expression profile in F1’ males (c), in genes having an over-expressed expression profile in F1’ females and in genes having an under-expressed expression profile in F1 females (e).

In genes following an intermediate profile in F1’ males we identified biological processes related to methylation, notably post-transcriptional modifications (eg., *RNA methylation,* Fig. 4a). Regarding genes over-expressed in F1 females compared to parental lines, biological processes over-represented were related to metabolic processes notably to integrate DNA, assemble organelles, or synthetize and transport of organic compounds (Fig. 4-b). DNA integration was also a process that was enriched in over-expressed genes in F1’ females (Fig. 4-d). Similarly, under-expressed genes in F1’ males were enriched in biological processes related to metabolic processes notably involved in ribosome assembly but also related to post-translational protein processing such as *protein folding* or *protein targeting the endoplasmic reticulum* (Fig. 4-c). Interestingly, some under-expressed F1’ males’ genes enriched in mitochondrial genes (which could suggest maternal effects). For F1 female under-expressed genes, only one biological process was enriched which was related to binding activity (Fig. 4-e). No functions expressly related to reproduction nor growth were found enriched (Supplementary file S2) and genes known to be involved in those functions (for instance reproduction) were rare and spread over the different gene expression profiles including intermediate (eg. Cathepsin B and serine/threonine phosphatase, Supplementary file S1: Table S1) and non-intermediate (eg., Cathepsin B-like, Cpeb genes, Supplementary file S1: Table S1) [66,67].

## Discussion

In this study, we aimed at describing first generation *S. haematobium* x *S. bovis* hybrids compared to their parents at the transcriptome level. Herein we provide an original reference transcriptome assembly allowing to map with similar rates *S. haematobium, S. bovis* and their reciprocal first-generation hybrids (F1 and F1’). We further quantified the number of genes falling into five main hybrid expression profiles namely over- and under-expressed, intermediate, *S. haematobium* like and *S. bovis* like expressed genes. Nearly half of the genes in hybrid progenies were differentially expressed when compared to the two parental lines, demonstrating the large gene expressions remodelling following hybridization. We showed that the most prevalent expression profile in *Schistosoma* hybrids was the intermediate profile, corresponding to gene expression levels in the hybrids that are significantly lower than the levels of expressions in one of the parental species while being significantly higher than the levels of expression observed in the other parental species. On the contrary, non-intermediate profiles (i.e., under-expressed, over-expressed and parent-like profiles) concerned a much smaller number of genes. Also, we detected very strong cross and sex effects on gene expression levels since genes falling into each expression profiles were very specific of each cross (F1 or F1’) and of each sex except for intermediate profiles. Finally, we showed that only few biological processes were characteristic of a given expression profile (eg., under-, over-expressed, parent-like or intermediate). Most enriched biological processes were in addition not directly related to traits potentially involved in hybrid vigour such as reproduction or growth, but were related to post-transcriptional regulation and to various metabolic processes and were inconsistent through sexes and crosses in each expression profile.

One of the main challenges of this study was to produce a transcriptome that can serve as a common reference for two sister species and their hybrids without being strongly biased toward one of the two parental species. There is currently no real consensus about how to analyse gene expression in hybrid species due to the difficulty to map hybrid RNA-seq reads to a single common reference [68]. Oftentimes strategies employed follow two main paths. On one hand, *de novo* approaches are often used because they allow to represent the genes of both parental species (eg., [6,69]) but the gene structure in introns and exons (for species for which a reference is available) is then lost on the way, along with the information about the species each genes is representative of, in addition chimeric transcripts can also be assembled. On the other hand, some studies chose to use the genome of one of the two parents to conserve genome structure information and ease analyses but oftentimes the mapping rate of the other species is lower, and the analysis of gene expression in hybrids is biased toward one of the parental species (eg., [5]). Here, we proposed a mid-way strategy relying on independent transcriptome assemblies of each parental species, based on their respective reference genomes, and a second step of identification of strictly orthologous genes. Our approach is promising because we obtained very similar mapping rates for the two parental species but also for their hybrids, hence gene expression of the hybrids can be assessed without being biased toward one of the two parental species. Also, we achieved a good completeness in term of metazoan orthologous genes retrieved. Although our approach could be improved (notably because we identified only very strict orthologous genes and so a *S. bovis* read could still map on a gene tagged as *S. haematobium* specific), it paves the way for further possibilities of transcriptome assemblies in the context of hybrid gene expression studies.

We found that up to 55% of genes were differentially expressed in *Schistosoma* hybrids compared to at least one of the parental species. First generation hybrids benefit from an entire new set of alleles brought from the combination of the two parental species. This “genomic shock” as proposed by Barbara McClintock [70] usually results in a disruption of gene expression regulation and in the activation of genetic mobile elements (and potential transposable elements), leading to new patterns of gene expression [71]. Here we did not find particular over-expression of transposable elements, but we did show that most of the genes in first generation hybrids present a new pattern of expression compared to at least one of the parental species. In particular the most prevalent gene expression profile we found was the intermediate one, which is an expression pattern that is significantly different from both parental species. Therefore, an important remodelling of the levels of gene expression is expected after first generation crosses and this was observed in our work.

We found that most hybrid genes were expressed at intermediate levels compared to their parental species. We expected that the more two genomes are compatible, the less conflict in expression will occur and the more likely intermediate expression profile would be. The predominance of intermediate expression profile in *S. haematobium x S. bovis* hybrids suggests that there are overall few conflicts between *S. haematobium* and *S. bovis* gene expression which result in most genes having an intermediate expression profile in first generation hybrids compared to their parents. This makes particular sense for our *Schistosoma* species since for instance *S. haematobium* and *S. bovis* are documented to present genomes that are highly permeable one to the other [24]. More generally, recent studies tend to demonstrate introgression to various levels (up to 80% in *S. haematobium x S. bovis* case) in many schistosomes species [23,38,72–74]. In this context the *S. haematobium* lineage from Cameroon we used in this study has recently been shown to be introgressed with *S. bovis* (see [75]). Similarly, at the transcriptomic level, mating between *S. haematobium* and *S. bovis* have been found to trigger only few differential gene expressions which also suggests that the two species remain quite co-adapted one to the other in term of gene regulation. We thus suggest that the abundant intermediate gene expression profiles we found in *S. haematobium x S. bovis* hybrids might be another facet of the signature of the high permeability that still exists between these two parasite species.

*Schistosoma* first generations hybrids are not only fertile, but may also have the particularity to present heterosis (higher infectivity, production of more numerous and larger eggs, [24,25,38,39]. Heterosis is commonly expected to result either from complementary expression of two inbred parental species (dominance model) or from synergetic interaction between parental alleles (overdominance model) [9]. In terms of gene expression, both models are thus expected to translate into either over-expressed, under-expressed or parent-like expression profile (i.e., non-intermediate expression profiles, [42]). In *Schistosoma* hybrids a total of 3,665 genes (22%) presented a non-intermediate expression profile, whereas almost double this number of genes (6,308, 38%) presented an intermediate profile (Fig. 2, Supplementary file S1: Table S1). Our results are hence quite new because non-intermediate profiles are usually found as the most prevalent profiles associated with hybrid vigour (eg., [4,5,69,76]). The dominance model relies on the assumption that hybrids perform better than inbred parental lines by expressing the parental allele that will be most advantageous at each gene, hence resulting in parent-like expression profiles [9,42]. However, we found quite a low prevalence of genes following a parent-like expression profile (5% of genes). We thus propose that if any, the over-dominant model should be privileged to explain *Schistosoma* hybrid vigor with a potential critical role of a small portion of genes (22%) that are over-expressed (and/or repressor genes that are under-expressed) since those profiles are more prevalent. Alternatively, we could also hypothesize that intermediate gene expression levels could actually be a form of dominance, if, within a gene, hybrids are over-expressing the parental allele that is advantageous while under-expressing the parental allele that are not advantageous. Such a pattern would also result in an intermediate expression profile at the gene level, but would require in order to be tested to have information about the parental origin of each allele and allele specific expression. This later hypothesis would however align with the general observation that *S. bovis* harbor an overall higher fitness on several traits (such as the size and number of eggs) and so do hybrids backcrossed with *S. bovis* [20]. It is important to bear in mind that intermediate expressed gene in first generation hybrids can still be over-expressed compared to the level of expression of this gene in *S. haematobium*, hence participating to the hybrid vigor observed on first generation hybrids.

Heterosis is a multigenic complex trait for which no consensus has yet been found. Some authors suggest that it may be brought by a cumulative effect of the differential expression of a variety of genes involved in one or several metabolic pathways affecting yield or energy use efficiency [77,78]. The few main biological processes we found enriched in hybrid expression profiles were related to post-transcriptional regulation (including RNA methylation) various metabolic processes (including DNA integration and organelle assembly), and mitochondrial gene expression (only for under-expressed expression profiles in F1’ males). Finding a process such as mitochondrial gene expression is interesting because it could reflect some sort of maternal effects since those are directly inherited from the mother and their genes differentially expressed in the offspring [79]. Similarly, heterosis could be shaped at least partly through post-transcriptomic regulation as our results are suggesting. However, the fact those processes were enriched inconsistently through crosses and sexes for each expression profile makes it difficult to draw general conclusion. Overall, our findings are rather unexpected because we did not detect processes involved in either growth or reproduction [42,80]. Some authors suggested that heterosis could alternatively result from a general dysregulation affecting all biological processes (rather than from a direct over expression of reproductive and growth processes) and that would indirectly increase (in an unbalanced way) the expression of reproduction and growth traits [7,10]. In that case, mis-regulated processes would not necessarily appear as enriched because they are covering the whole range of biological processes expressed. This scenario would match better our observations and especially the fact that genes involved in each profile were quite cross and sex-specific. We thus suggest that i) heterosis in *S. haematobium x S. bovis* hybrids might be shaped at least partly through post-transcriptional regulation and involving various metabolic processes, but ii) that more likely a general mis-regulation of main biological processes indirectly benefit hybrid worm growth and reproduction notably in first generation hybrids.

It is worth noting that for a long-time hybrid vigor has been studied independently from hybrid incompatibility. As such, studies aiming at identifying the molecular basis of hybrid vigor are still often restricted to model species or species of agroeconomic interest [4–6,69]. However, some studies focusing on the molecular basis of hybrid incompatibilities have investigated wild species [81,82]. Interestingly both type of studies highlighted the presence of genes having over-expression profiles but gave them a very contrasted interpretation (base of hybrid vigor on one hand and proof of mis-regulation in the other hand (eg., [6,82]). This observation along with our results seem to support the idea that hybrid vigor is one facet of hybrid incompatibility. Here our findings can be interpreted at the light of the idea exposed in [7] stating that hybrid vigor can result from a mis-regulation of multiple process, preferably involved in trade-off with reproduction and growth functions. We hypothesize that when hybridizing, *S. haematobium* and *S. bovis*, random genes (explaining why those are different between hybrid crosses and sexes) interact at the first hybrid generation and are expressed following one of the hybrid expression profiles we described. Those genes do not affect one specific biological process but rather all of them at a smaller scale. This general mis-regulation could result in hybrid vigor in the first generation as the expression of an unbalanced phenotype which is oftentimes not maintained over generations [7]. Our hypothesis could explain why even when pre-zygotic barriers are overcome, first-generation *S. haematobium x S. bovis* hybrids remain rare in the wild (even though the hypothesis that *S. haematobium* and *S. bovis* do not meet often due to their final host species and tropism difference likely also contribute to this observation [23,83,84]). Indeed, the phenotype of the first-generation hybrids might not be sustainable long term unless a particularly fitted combination of gene occur. We nonetheless do not exclude that a rare favorable interaction between parental genes could have occurred in the past, giving birth to the current level of introgression documented in *Schistosoma* species [38,75,85].

In conclusion, in the current study we propose through an original approach one of the first detailed description of hybrid expression profiles in a non-model species (but see [19]). We showed that all types of hybrid expression profiles were found (over- and under-expressed, intermediate, and similar to one or the other parent) but that the far most abundant profile is the intermediate one. We suggest that a predominance of intermediate profiles could be the signature of parental species permeability rather than the basis of heterosis. Genes falling into the profiles of expression we detailed were dependent of the sex and on the direction of the hybrid cross and inconsistently coded for specific biological processes. We hence strongly suggest that hybridization between *Schistosoma* species leads to a general mis-regulation of genes coding for general molecular processes potentially given raise to hybrid vigor. We hence argue in favor of further investigations of gene interactions in wild hybrids to further assess how a mis-regulation of gene expression could promote heterosis in a hybrid phenotype. More generally we argue that more studies focusing on the genomic determinants of hybrid vigor in relation to potentially well-identified over-fitted phenotypes in *Schistosoma* hybrids are warranted.

## Acknowledgements

This work has been funded by the French Research National Agency (project HySWARM, grant no. ANR-18-CE35-0001). This study is set within the framework of the “Laboratoires d’Excellence (LABEX)” TULIP (ANR-10-LABX-41).

## Availability of data and materials

Raw sequences used in this study have been submitted to the Sequence Read Archive under the BioProject PRJNA491632. Further data and scripts necessary to replicate the study along with the transcriptome assembled and gtf annotation will be released upon publication in a figshare repository currently on restricted access for review purposes at https://figshare.com/s/a4730c7301a7f2961042.

## Competing interest statement

We have no competing interest.

## Supplementary material captions

**Supplementary file S1:**

**Fig. S1:** Principal Component Analysis run on gene expression samples of *S. haematobium, S. bovis* and their reciprocal F1 and F1’ hybrids.
**Fig. S2:** Heatmaps illustrating the gene clustering within each hybrid expression profiles in F1’ females (F1’) compared to *S. haematobium* (SH) and *S. bovis* (SB) females: under and over-expressed profiles (a), intermediate profiles (b), *S. haematobium* like profile (c) and *S. bovis* like profile (d).
**Fig S3:** Heatmaps illustrating the gene clustering within each hybrid expression profiles in F1 males (F1) compared to *S. haematobium* (SH) and *S. bovis* (SB) females: under and over-expressed profiles (a), intermediate profiles (b), *S. haematobium* like profile (c) and *S. bovis* like profile (d).
**Fig S4:** Heatmaps illustrating the gene clustering within each hybrid expression profiles in F1’ males (F1’) compared to *S. haematobium* (SH) and *S. bovis* (SB) females: under and over-expressed profiles (a), intermediate profiles (b), *S. haematobium* like profile (c) and *S. bovis* like profile (d).
**Fig. S5:** Venn plots showing shared genes involved in each hybrid expression profiles between the two hybrid crosses (F1 and F1’) and the two sexes.
**Table S1:** Table summarising the mapping rates of the *S. haematobium, S. bovis* and hybrids samples on the transcriptome assembled in this study.

**Supplementary file S2**: Table summarising for each gene of the assembled transcriptome between *S. haematobium* and *S. bovis*, their expression profile (if any), the cross and sex in which their expression profile has been measured, relevant log fold changes for contrasts between hybrids and parents, and their associated annotations.

**Supplementary file S3**: Table summarising the result of the gene ontology analysis and giving for each enriched Gene Ontology, their statistical significance (FDR and p-value), the number of genes involved in the Gene Ontology category and the gene expression profile of the gene set (*m* stands for male and *f* for female, *F1p* stands for F1’). The column type refers to the origin of the set of gene tested which can be either the differentially expressed gene of a particular hybrid profile (gene_set) or the rest of the genes in the reference transcriptome but not presenting a particular hybrid expression profile.

## Notes

### Competing Interest Statement

The authors have declared no competing interest.

